# Mediterranean Diet, Stress Resilience, and Aging in Nonhuman Primates

**DOI:** 10.1101/2020.09.25.313825

**Authors:** Carol A. Shively, Susan E. Appt, Haiying Chen, Stephen M. Day, Brett M. Frye, Hossam A. Shaltout, Marnie G. Silverstein-Metzler, Noah Snyder-Mackler, Beth Uberseder, Mara Z. Vitolins, Thomas C. Register

**Affiliations:** Department of Pathology/Comparative Medicine, Wake Forest School of Medicine; Department of Biostatistics and Data Science, Wake Forest School of Medicine; Department of Internal Medicine/Gerontology and Geriatric Medicine, Wake Forest School of Medicine; Department of Obstetrics and Gynecology, Wake Forest School of Medicine; School of Life Sciences, Center for Evolution and Medicine, Arizona State University; Department of Epidemiology & Prevention, Wake Forest School of Medicine

**Keywords:** Mediterranean diet, Stress Resilience, Hypothalamic-Pituitary-Adrenal, Autonomic Nervous System, Aging, Nonhuman Primates

## Abstract

Persistent psychological stress increases the risk of many chronic diseases of aging. Little progress has been made to effectively reduce stress responses or mitigate stress effects suggesting a need for better understanding of factors that influence stress responses. Limited evidence suggests that diet may be a factor in modifying the effects of stress. However, long-term studies of diet effects on stress reactive systems are not available, and controlled randomized clinical trials are difficult and costly. Here we report the outcomes of a controlled, randomized preclinical trial of the effects of long-term consumption (31 months, ∼ equivalent to 9 human years) of Western versus Mediterranean - like diets on behavioral and physiological responses to acute (brief social separation) and chronic (social subordination) psychosocial stress in 38 adult, socially-housed, female cynomolgus macaques. Compared to animals fed a Western diet, those fed the Mediterranean diet exhibited enhanced stress resilience as indicated by lower sympathetic activity, brisker and more overt heart rate responses to acute stress, more rapid recovery, and lower cortisol responses to acute psychological stress and adrenocorticotropin (ACTH) challenge. Furthermore, age-related increases in sympathetic activity and cortisol responses to stress were delayed by the Mediterranean diet. Population level diet modification in humans has been shown to be feasible. Our findings suggest that population-wide adoption of a Mediterranean-like diet pattern may provide a cost-effective intervention on psychological stress and promote healthy aging with the potential for widespread efficacy.

**Highlights:**

- There is no population level treatment to reduce stress and associated disease.
- Mediterranean diet reduced sympathetic activity.
- Mediterranean diet reduced cortisol response to acute stress and to ACTH challenge.
- Mediterranean diet delayed age-related increases in sympathetic activity and cortisol responses to stress.
- These results suggest a dietary strategy to increase stress resilience.

## Introduction

Psychological stress is an important determinant of physiology, and may cause adverse health outcomes. Americans report some of the highest perceived stress levels in the world (1). Chronic psychological stress increases the risk of many prevalent diseases including depression and anxiety, as well as diseases of aging such as obesity, metabolic syndrome, type 2 diabetes, cardiovascular disease, stroke, and Alzheimer’s Disease (2-8). Attempts to reduce stress levels are often ineffective because stressors are uncontrollable, and effective therapeutic interventions are difficult to adhere to or unavailable due to financial constraint. Attempts to intervene on the psychological stress-disease relationship thus have not been successful, suggesting a lack of understanding of factors that may contribute to this relationship.

Adverse health effects of psychological stress may be amplified by other environmental stressors including an unhealthy diet. Many Americans consume a Western diet rich in animal protein and saturated fat, salt, and sugar which increases disease risk. Observational studies suggest lower perceived stress is associated with high fruit, vegetable, and protein intake (9-11) and adherence to a Mediterranean diet pattern (12-14). Likewise, associations have been reported between higher cortisol levels and high simple sugar and saturated fat intake (9, 15). Small, short-term, crossover studies in humans suggest that high fat diets, particularly those rich in saturated animal fats, increase cardiovascular responses to a standardized stressor (16-18). Similarly, small, short-term rodent studies suggest that high saturated fat feeding may increase hypothalamic-pituitary-adrenal (HPA) responses to stressors (19, 20). Thus, the limited clinical and preclinical data available are consistent with the hypothesis that diet composition may affect physiological stress reactivity. Whether these responses are sustained chronically is unknown.

Limited data from nonhuman primate (NHP) studies are also consistent with the hypothesis that diet may exacerbate physiological responses to stressors, particularly social subordination. Social subordination is stressful in female macaques (*Macaca spp.*) as they receive more aggression, less affiliative attention, spend more time alone, and have impaired ovarian function, heightened heart rate (HR) responses to acute stressors, and high circulating cortisol concentrations relative to their dominant counterparts (21). Subordinates also have lower bone density, greater visceral obesity, more inflammation (22), increased biomarkers of breast and endometrial cancer risk, are more likely to exhibit depression-like phenotypes, and develop more diet-induced coronary and carotid artery atherosclerosis than their dominant counterparts (23-25). Notably, all of these observations were made in macaques consuming a Western-like diet.

HPA axis and autonomic nervous system (ANS) function have been characterized in female macaques consuming monkey chow (the standard diet fed to captive monkeys) in some studies and Western-like diets in other studies. Intriguingly, post hoc comparisons suggest that the cortisol response to adrenocorticotropin (ACTH) is higher in subordinate than dominant macaques in studies in which animals are fed a Western-like diet (26, 27), whereas the cortisol response to ACTH is lower in subordinates than dominants in studies in which monkey chow is fed (28). Similarly, compared to baseline measured during monkey chow consumption, 24 hour heart rates increased in subordinates, but not dominants, after long term Western diet consumption (29). The differences in effects may be due to differences in diets. The Western diet in these studies was modeled on the typical North American diet and contained 40% of calories from fat (mostly saturated), 0.25-0.40 mg cholesterol/kcal (450-700 mg cholesterol /day human equivalent), with protein and fat primarily derived from animal sources. In contrast, monkey chow is very low in fat (13% of calories) and cholesterol (trace amounts), with protein and fat almost entirely from vegetable sources (Diet #5037/5038, LabDiet, St. Louis, MO). The translational relevance to human health of physiological responses when monkey chow is fed is unknown since monkey chow has no human diet parallel. (see Table 1 for chemical composition comparison of human and nonhuman primate diets.) Thus, a more human-relevant diet comparison is needed.

**Table 1.**
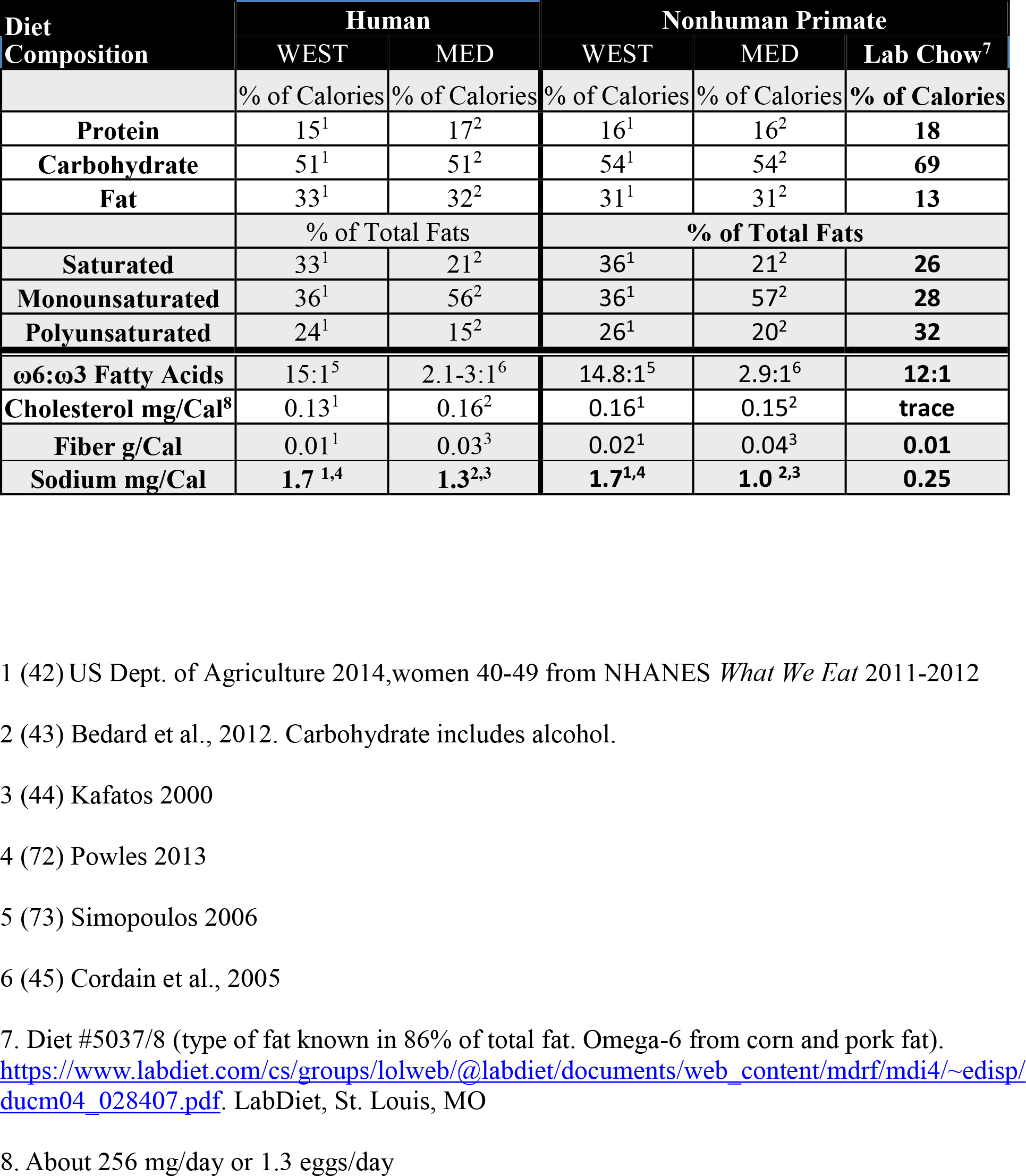
Macronutrient Content of Experimental Diets and Monkey Chow.

| Diet Composition | Human |  | Nonhuman Primate |  |  |
| --- | --- | --- | --- | --- | --- |
|  | WEST | MED | WEST | MED | Lab Chow<sup>7</sup> |
|  | % of Calories | % of Calories | % of Calories | % of Calories | % of Calories |
| **Protein** | 15<sup>1</sup> | 17<sup>2</sup> | 16<sup>1</sup> | 16<sup>2</sup> | **18** |
| **Carbohydrate** | 51<sup>1</sup> | 51<sup>2</sup> | 54<sup>1</sup> | 54<sup>2</sup> | **69** |
| **Fat** | 33<sup>1</sup> | 32<sup>2</sup> | 31<sup>1</sup> | 31<sup>2</sup> | **13** |
|  | % of Total Fats |  | % of Total Fats |  |  |
| **Saturated** | 33<sup>1</sup> | 21<sup>2</sup> | 36<sup>1</sup> | 21<sup>2</sup> | **26** |
| **Monounsaturated** | 36<sup>1</sup> | 56<sup>2</sup> | 36<sup>1</sup> | 57<sup>2</sup> | **28** |
| **Polyunsaturated** | 24<sup>1</sup> | 15<sup>2</sup> | 26<sup>1</sup> | 20<sup>2</sup> | **32** |
| **ω6:ω3 Fatty Acids** | 15:1<sup>5</sup> | 2.1-3:1<sup>6</sup> | 14.8:1<sup>5</sup> | 2.9:1<sup>6</sup> | **12:1** |
| **Cholesterol mg/Cal<sup>8</sup>** | 0.13<sup>1</sup> | 0.16<sup>2</sup> | 0.16<sup>1</sup> | 0.15<sup>2</sup> | **trace** |
| **Fiber g/Cal** | 0.01<sup>1</sup> | 0.03<sup>3</sup> | 0.02<sup>1</sup> | 0.04<sup>3</sup> | **0.01** |
| **Sodium mg/Cal** | **1.7**<sup>1,4</sup> | **1.3**<sup>2,3</sup> | **1.7**<sup>1,4</sup> | **1.0**<sup>2,3</sup> | **0.25** |

1 (42) US Dept. of Agriculture 2014, women 40-49 from NHANES *What We Eat* 2011-2012
2 (43) Bedard et al., 2012. Carbohydrate includes alcohol.
3 (44) Kafatos 2000
4 (72) Powles 2013
5 (73) Simopoulos 2006
6 (45) Cordain et al., 2005
7. Diet #5037/8 (type of fat known in 86% of total fat. Omega-6 from corn and pork fat). https://www.labdiet.com/cs/groups/lolweb/@labdiet/documents/web_content/mdrf/mdi4/~edisp/ducum04_028407.pdf. LabDiet, St. Louis, MO
8. About 256 mg/day or 1.3 eggs/day

Controlled clinical trials of long-lasting changes in stress responses due to diet are difficult and costly. Human population studies are not definitive since dietary data and stress levels are self-reported, often collected retrospectively, and nutrient intakes are drawn from food composition tables based on self-reports. Furthermore, observational studies compare individuals who <u>choose</u> to consume either a prudent or Western diet, and these two groups may differ in other characteristics that could influence health such as socioeconomic status (SES), alcohol consumption, smoking behavior, and exercise levels (30, 31). Finally, other factors that affect stress reactivity (e.g. physical activity, medications) may be inaccurately reported, and in cross-sectional studies, there is the possibility of reverse causality (i.e. stressors may influence food choices).

Thus, we conducted a longitudinal, randomized preclinical trial in female cynomolgus macaques (*Macaca fascicularis*) to test the hypothesis that long term Western (WEST), compared to Mediterranean (MED), diet consumption would result in greater physiological stress reactivity. We examined the effects of diet on the ANS and the HPA axis because they have bidirectional effects on each other, and independent and synergistic effects on molecular and cellular processes that promote downstream pathologies when chronically activated (2, 7, 32-34). We focused on females because of mammalian sex differences in stress responses (35), and because much of the literature on social status differences in stress responses and their relationship to health outcomes on which this study builds is from female NHPs. As the lifespan of these NHPs is approximately 3-4 times shorter than humans, the 31-month treatment period is approximately equivalent to 9 human years. Thus, changes over time may reflect aging effects as the NHPs entered the study at middle-age. Important to this investigation, stress resiliency is thought to slow aging and improve overall health (36). We chose to study diet patterns rather than single dietary constituents because human diets include multiple characteristics that may affect health and be synergistic (37). The experimental diets were formulated to emulate dietary patterns consumed by humans. The NHPs were socially housed, and the study focused on responses to the chronic stress of social subordination, and to the acute stress of brief social separation, which are commonly experienced by this species. Previously, we reported that subjects in the WEST group ate more, developed greater adiposity, insulin resistance, and hepatosteatosis, had different gut microbiome populations, and exhibited higher pro-inflammatory gene expression in monocytes relative to those that consumed the MED diet (38, 39). Here we demonstrate that, compared to the WEST diet, the MED diet promotes stress resilience as evidenced by greater sensitivity to and rebound from acute stressors, and reduced physiological responses to chronic stressors.

## Materials and Methods

### Subjects

Forty-three adult, middle-aged (X=9.0, range=8.2-10.4 years, estimated by dentition) female cynomolgus macaques were obtained (SNBL USA SRC, Alice, TX), quarantined in single cages for one month, then moved to social groups of four animals each which lived in 3m X 3m X 3m enclosures with daylight exposure, on a 12/12 light/dark cycle (light phase: 0600-1800), with monkey chow and water available *ad libitum*.

### Experimental Design

The monkeys habituated to their social groups and were characterized during a 7-month Baseline phase while consuming monkey chow (diet 5037/5038, LabDiet, St. Louis, MO; **Table 1**). The monkeys were randomly assigned to the WEST or the MED diet group for 31 months (the equivalent of about 9 human years; **Supplementary Table 1**). The two groups were confirmed to be balanced on pretreatment characteristics that reflected overall health including body weight, body mass index, age, basal cortisol concentration, and plasma lipid concentrations measured as previously described (40, 41). All animal manipulations were performed in accordance with state and federal laws, the US Department of Health and Human Services, the National Institutes of Health guide for the care and use of Laboratory animals, and the Animal Care and Use Committee of Wake Forest School of Medicine.

### Experimental Diets

The experimental diets were designed to mimic human Western or Mediterranean diet patterns (see diet compositions in **Supplementary Table 2)**. These semi-purified diets were formulated to be isocaloric with respect to protein, fat and carbohydrate macronutrients, and identical in cholesterol content (∼ 320mg / 2000 kilocalories (Cals)/day) (**Table 1**). The WEST diet was formulated to be similar to that consumed by American women age 40-49 as reported by the US Department Agriculture (42). The diet contained protein and fat derived mainly from animal sources, was high in saturated fats and sodium, and low in monounsaturated and n-3 fatty acids. The MED diet was formulated to mimic key aspects of the MED diet with protein and fats derived mainly from plant sources, some lean protein from fish and dairy, and a relatively high monounsaturated fatty acid content (43, 44). The MED diet n-6:n-3 fatty acid ratio was similar to a traditional hunter-gatherer type diet (45), was high in complex carbohydrates and fiber, and lower than the WEST diet in sodium and refined sugars. Key MED ingredients included English walnut powder and extra-virgin olive oil which were the primary components provided to participants in the PREDIMED study (46). Both test diets differ considerably from monkey chow (**Table 1; Supplementary Note 1**).

### Food Consumption

To measure consumption, monkeys were trained to enter single cages that were placed inside their social group enclosures. They were fed for two hours between 0700 and 0900, and then released back into the social group enclosure. Because nearly all consumption occurred in 30 min or less, at the beginning of Experimental month 12, the morning feeding was shortened to 1 hour (0700-0800) and an additional feeding was added in the afternoon (1500-1600).

### Behavior

Behavior was recorded during 10 minute focal observations (47) twice/week for 6 weeks during the Baseline phase after 3 months of social grouping (2 hours/animal), and for 24 months during the Experimental phase (37 hours/animal). As in previous studies (21, 48), levels of social stress were assessed as the frequency a monkey was aggressed by another monkey, how much time was spent fearfully scanning the social group (i.e. head swiveling rapidly while grimacing and lip-smacking), and how much time a monkey was groomed by another monkey (which is inversely related to social stress). Baseline behavioral observations were only used to determine social status. Social status was determined by which monkey submitted in agonistic interactions and was stable over time; only 3/40 (7.5%) animals changed rank over the 38-month study period (Spearman’s rho Baseline • Experimental phase ranks=0.90, p<0.0001). For all analyses, those in the top half of the hierarchy in their social group were considered dominant, and those in the bottom half were considered subordinate.

### Blood Pressure

Blood pressure was measured during the Baseline phase (6 months after social grouping) and 26 months after experimental diet consumption, at about the same time of day (0800). Following sedation (ketamine HCl 15 mg/kg) and stabilization of HR, systolic (SBP), diastolic (DBP), and mean arterial pressure (MAP), measurements were taken using high definition oscillometry via tail cuff.

### 24 Hour HR via Telemetry

HRs were recorded during the Baseline phase (3 months after social grouping) and after 12 and 29 months of experimental diet consumption. Monkeys were captured, sedated with ketamine HCl (10 mg/kg), and outfitted with a nylon mesh protective jacket over a portable electrocardiogram telemetry unit (Life Sensing Instrument Co, Tullahoma, TN, US). After overnight recovery from sedation, HR recording began and continued for 24 hours. Interbeat intervals were recorded at a sampling rate of 250 Hz. HR hourly averages were calculated for analysis (41, 49, 50).

### Heart Rate Variability (HRV)

HRV was assessed in 2-hour nighttime (0100-0300) and daytime (1600-1800) blocks from the 24-hour telemetry data. The nighttime block represents the nadir of the circadian HR rhythm. The daytime block was chosen because it was the only 2-hour block during lights on in which no animal care and research activities were done. Heart rate recordings were analyzed using Nevrokard-HRV software (Nevrokard Kiauta, d.o.o. Izola Slovenia) to obtain the HRV measures. HRV was assessed as the standard deviation of successive interbeat intervals from which artifacts have been removed (SDNN), and the root mean square of successive interbeat intervals measured between the R-to-R peaks (RMSSD). Power spectral analysis with Fast Fourier transformation was used to determine the very low frequency (%VLF) component (range: <0.01Hz) which represents activation of the sympathetic nervous system (SNS), the low frequency (%LF) component (range: 0.01-0.2Hz) which represents the activity of both the parasympathetic (PNS) and SNS, the high frequency (%HF) component (range: 0.2-0.8Hz) which represents activity of the parasympathetic nervous system (PNS), and the ratio of LF/HF, a measure of sympathovagal balance (51, 52).

### Social Separation Test of Responses to Acute Stress

This test was conducted during the Baseline phase (4 months after social grouping) and after 12 and 29 months of experimental diet consumption, using a procedure adapted from Michopoulos et al (2012; diagrammed in **Figure 4)**. Briefly, the animals were habituated to moving into single cages on voice command and to allow trained technicians to draw a blood sample during months 2 and 3 of the Baseline phase. Twenty-six hours prior to the acute stress test the monkeys were outfitted with electrocardiography telemetry devices. HR was recorded continuously beginning 2 hours prior to the onset of the stress test and for 4 hours thereafter. On test day at 0550 (10 min prior to lights on), the monkey exited the home cage on voice command, entered a single cage, and was moved to a room where a baseline blood sample was taken before adrenal cortisol increases would be detectable, less than 9 min after the technician entered the monkey building (baseline sample). Blood sampling was repeated at 30 min (stress sample) after which the monkey was returned to her home enclosure. This procedure was repeated, and blood samples taken at 120 and 240 min (recovery samples). Cortisol concentrations were determined in all blood samples.

### Cortisol Response to ACTH challenge

This test, which measures adrenocortical secretion of cortisol in response to exogenous ACTH independent of hypothalamic–pituitary activity, was done during the Baseline phase (5 months after social grouping), and after 13 and 30 months of experimental diet consumption as previously described (50). Following an overnight fast, the animals were administered dexamethasone (0.5 mg/kg) to suppress hypothalamic–pituitary activity. Four hours later they were sedated (ketamine hydrochloride 15 mg/kg), and a blood sample was taken (<9 min from entering the monkey building) for the determination of baseline cortisol. The animals were administered the ACTH challenge (Cortrosyn^®^, Organon, Inc., 10 ng/kg IV), blood samples were taken 15 and 30 min later for cortisol measurement (26), and the area under the cortisol curve was calculated (50).

### Cortisol Assay

Acute stress test Baseline phase cortisol samples were assayed with radioimmunoassay (RIA) kits from Diagnostic Products Corporation (Los Angeles, California). Due to discontinuation of all DPC RIAs by Siemens, Experimental phase samples were assayed using RIA kits from DiaSource (IBL America). All samples were run within a single assay lot number for the respective platforms. Intraassay CVs within the platforms were <7% for controls and pools. Validation studies between platforms were conducted in which samples which had previously been assayed on the DPC platform were then subjected to analyses using RIAs from IBL. Linear correlations were observed between the different platforms (r>0.8), although the slope of the line was not =1. Two separate IBL kits provided highly correlated values (r=0.97). In order to align the DPC and IBL assays, data from the DPC assay were recalculated for alignment with the IBL assay to insure continuity across the study.

### Data Analysis

The primary analyses were performed using 2 _dominant, subordinate_ X 2 _WEST, MED_ analyses of variance (ANOVA), or covariance (ANCOVA) where indicated, with repeated measures for dependent variables measured at multiple time points. HRV analyses included a term for time of day. Follow up comparisons were made with post hoc Tukey tests. Data were transformed as needed to improve homogeneity of variance. All graphed data are the means and standard errors of the raw data. ANCOVA was used when Baseline measures accounted for a significant proportion of the variance, and means were adjusted for Baseline where indicated. To simplify the model, heart rate data were analyzed at each time point (baseline, 12, and 29 months) separately. Cortisol area under the curve (AUC) was calculated using the integration method. The significance level was set at p=0.05 and all reported p values are two-sided. Effect sizes are reported as mean and per cent differences for key effects of diet.

## Results

A total of 38 animals are included in the analysis (N_WEST_=21, 11 dominant, 10 subordinate; N_MED_=17, 10 dominant, 7 subordinate). Two animals did not tolerate the experimental diets, were fed standard monkey chow and excluded from analyses reported here, and three animals died during the study (**Supplementary Note 2**).

### 1. Behavior (Figure 1)

During the Experimental phase socially subordinate females received more aggression (F[1, 34]=22.2, p<0.00001), were groomed less (F[1, 34]=5.33, p=0.027), and spent more time fearfully scanning their enclosure (F[1, 34]=6.62, p=0.015) than their dominant counterparts. There were no significant effects of diet or interactions between diet and social status interactions on these behaviors (all p’s>0.10).

### 2. Blood Pressure (Figure 2)

#### 2.a. Diet and Social Status Effects

There were no main or interaction effects of diet on BP (all p’s>0.10). At Baseline, subordinates had higher SBP than dominants (F[1, 37]=5.75, p=0.02; **Figure 2a**). Over the course of the study SBP decreased in subordinates and increased in dominants converging at 26 months (Status X Time F[1, 33]=4.67, p=0.038; **Figure 2a**).

**Figure 1.**
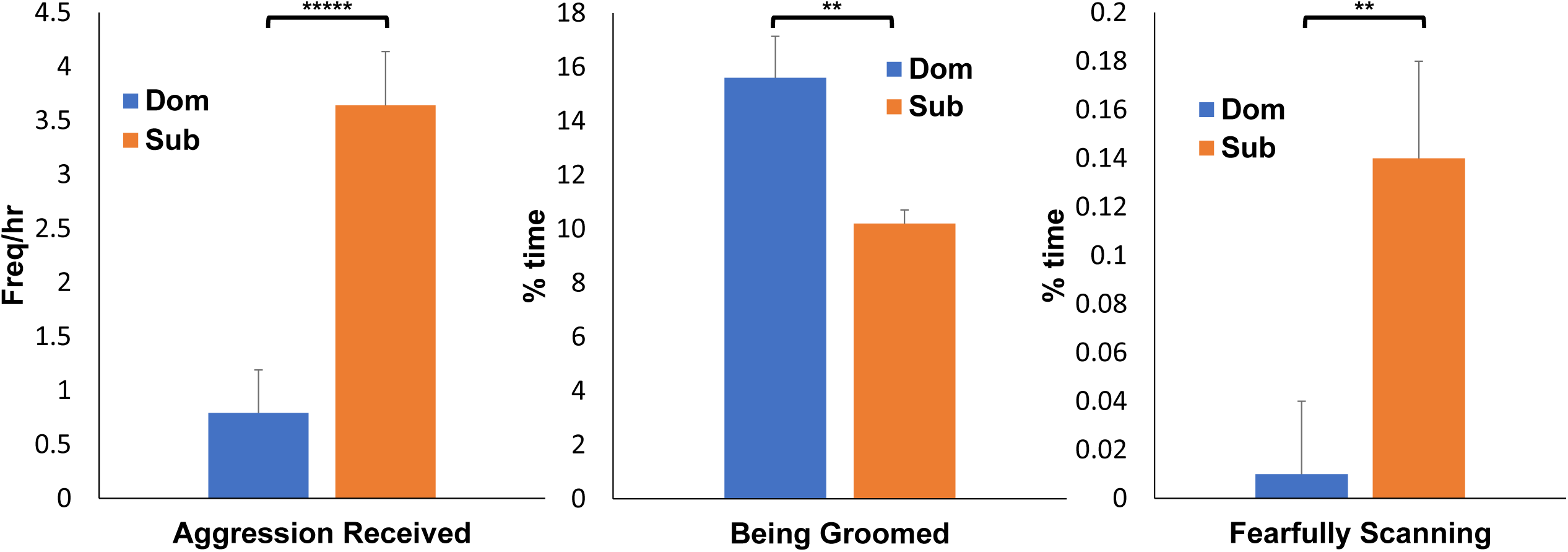
Social Status Differences in Stress-Related Behaviors. During the Experimental phase socially subordinate females received more aggression (F[1, 34]=22.2, p<0.00001*****), were groomed less (F[1, 34]=5.33, p=0.027**), and spent more time fearfully scanning their enclosure (F[1, 34]=6.62, p=0.015**) than their dominant counterparts. There were no significant effects of diet or diet by social status interactions on these behaviors (all p’s>0.10).

**Figure 2.**
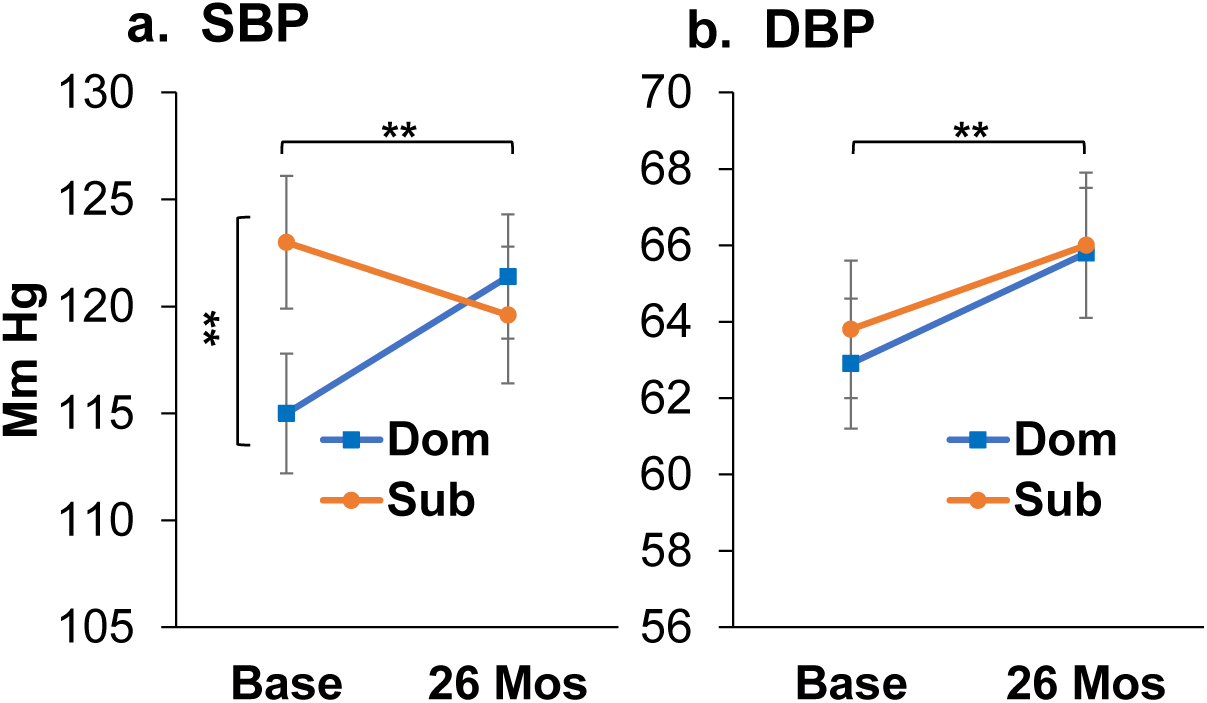
Social Status and Aging Effects on Blood Pressure: **a**. At Baseline, subordinates had higher SBP than dominants (F[1,37]=5.75, p=0.02**); Subordinates decreased, and dominants increased SBP after 26 months (Status X Time F[1, 33]=4.67, p=0.038**); **b**. Diastolic blood pressure (DBP) increased over the course of the experiment (main effect of Time F[1, 33]=5.03, p=0.03**).

#### 2.b. Aging Effects

DBP increased in all animals between Baseline and 26 months by 2.5 mmHg on average (Time F[1, 33]=5.03, p=0.03, **Figure 2b**; **Supplementary Note 3**).

### 3. 24 Hour Heart Rates (Figure 3)

#### 3.a. Diet and Social Status Effects

Each time point (Baseline, 12, and 29 months) was examined individually. There were no main or interaction effects of diet or social status at Baseline or after 12 months of diet consumption (ANCOVA controlling for baseline), and no main or interaction effects of social status at 29 months (ANCOVA controlling for baseline) (all p’s>0.22; **Supplementary Figure 1)**. At 29 months there was a significant diet X hour interaction (ANCOVA controlling for baseline F[23, 766]=2.5, p=0.0001; **Figure 3a**). HRs of monkeys consuming the MED diet spiked sharply at mealtimes, and recovered between meals, whereas peaks and recoveries were less pronounced and more sluggish in the WEST group (0700 vs 1100 hours: MED Tukey p<0.0001; WEST Tukey p=0.264; differences between adjusted means: MED=17 bpm; WEST=3 bpm).

**Figure 3.**
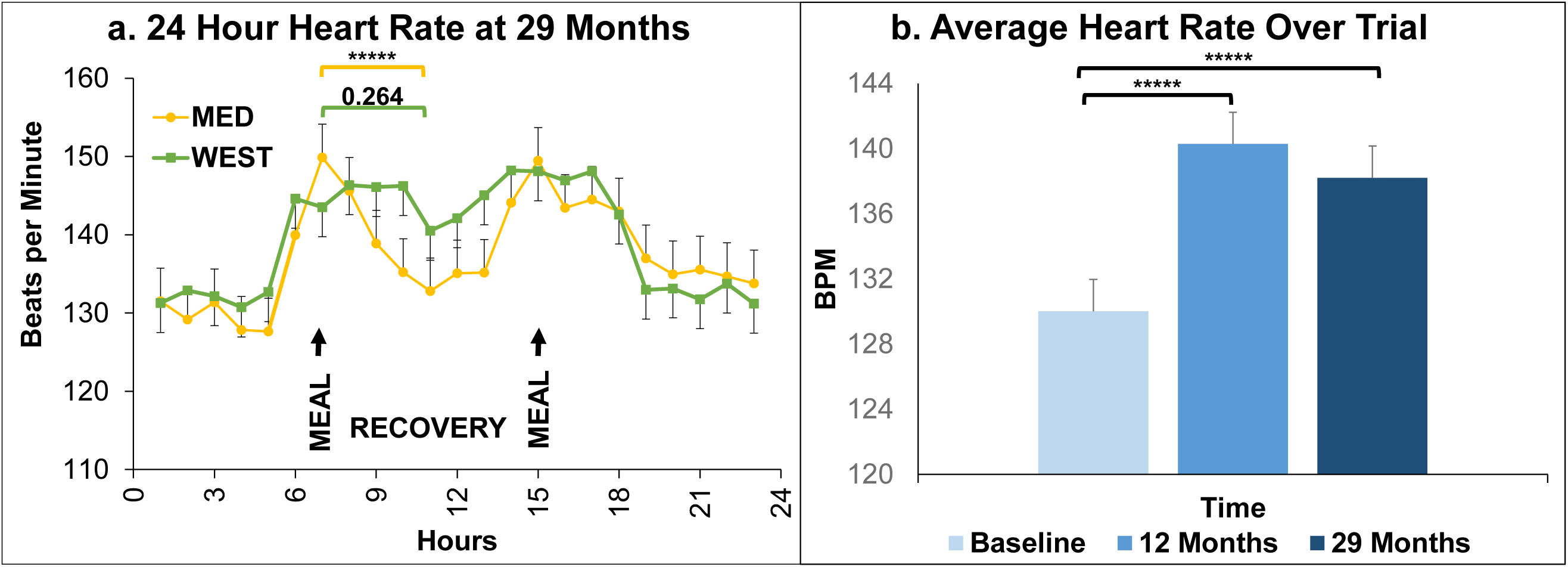
24 Hour Heart Rates. Diet and Social Status Effects: Baseline, 12-month, and 29-month time points were examined individually. There were no main or interaction effects of diet or social status at Baseline or after 12 months of diet consumption, and no main or interaction effects of social status at 29 months (all p’s>0.22; **Supplementary Figure 1). a**. At 29 months there was a significant diet X time interaction (ANCOVA controlling for baseline F[23, 766]=2.5, p=0.0001). The HRs of monkeys consuming the MED diet spiked sharply at mealtimes, and recovered significantly in between meals, whereas peaks and recoveries were less pronounced and more sluggish in the WEST group (0700 vs 1100 hours: MED Tukey p<0.0001*****; WEST Tukey p=0.264). **b. Aging Effects:** Heart rates, averaged over 24 hours, increased over the course of the experiment (3 time points X 2 social status X 2 diets X 24 hours ANOVA; main effect of experimental time point: F[2, 2397]=169.8, p<0.0001). Overall, HRs increased between Baseline and 12 months (Tukey p<0.0001*****) and between Baseline and 29 months (Tukey p<0.0001*****).

#### 3.b. Aging Effects

Average heart rate over 24 hours increased over the course of the experiment (3 time points X 2 social status X 2 diets X 24 hours ANOVA; main effect of experimental time point: F[2, 2397]=169.8, p<0.0001; **Figure 3b**). Overall, HRs increased between Baseline and 12 months (Tukey p<0.0001) and between Baseline and 29 months (Tukey p<0.0001; **Supplementary Figure 1**).

### 4. Heart Rate Variability During 24-hour HR Recording

#### 4.a. Diet and Social Status Effects (Figure 4)

At Baseline there were no main or interaction effects of status, diet, or time of day for any HRV parameter (all p’s>0.09). Furthermore, Baseline values of HRV parameters did not account for a significant proportion of the variance in ANCOVA, therefore repeated measures ANOVA was used to examine the main and interaction effects of status and diet during the Experimental phase. Dominants had higher heart rate variability (SDNN (F[1, 34]=4.685, p=0.038) and RMSSD (F[1, 34]=6.526, p=0.015) than subordinates (**Figure 4a**). Compared to the WEST group, the MED diet group had 2.7% higher %LF (F[1, 34]=4.322, p=0.045; mean difference=1.54). %VLF tended to be lower in MED animals, but this was not statistically significant (F[1, 34]=3.453, p=0.072; **Figure 4b**). These observations suggest that animals consuming the MED diet had lower sympathetic activity overall than the WEST group. (See **Supplementary Table 1** for raw means and standard errors.) There were no other main or interaction effects of diet and social status at any time point (all p’s>0.10).

**Figure 4.**
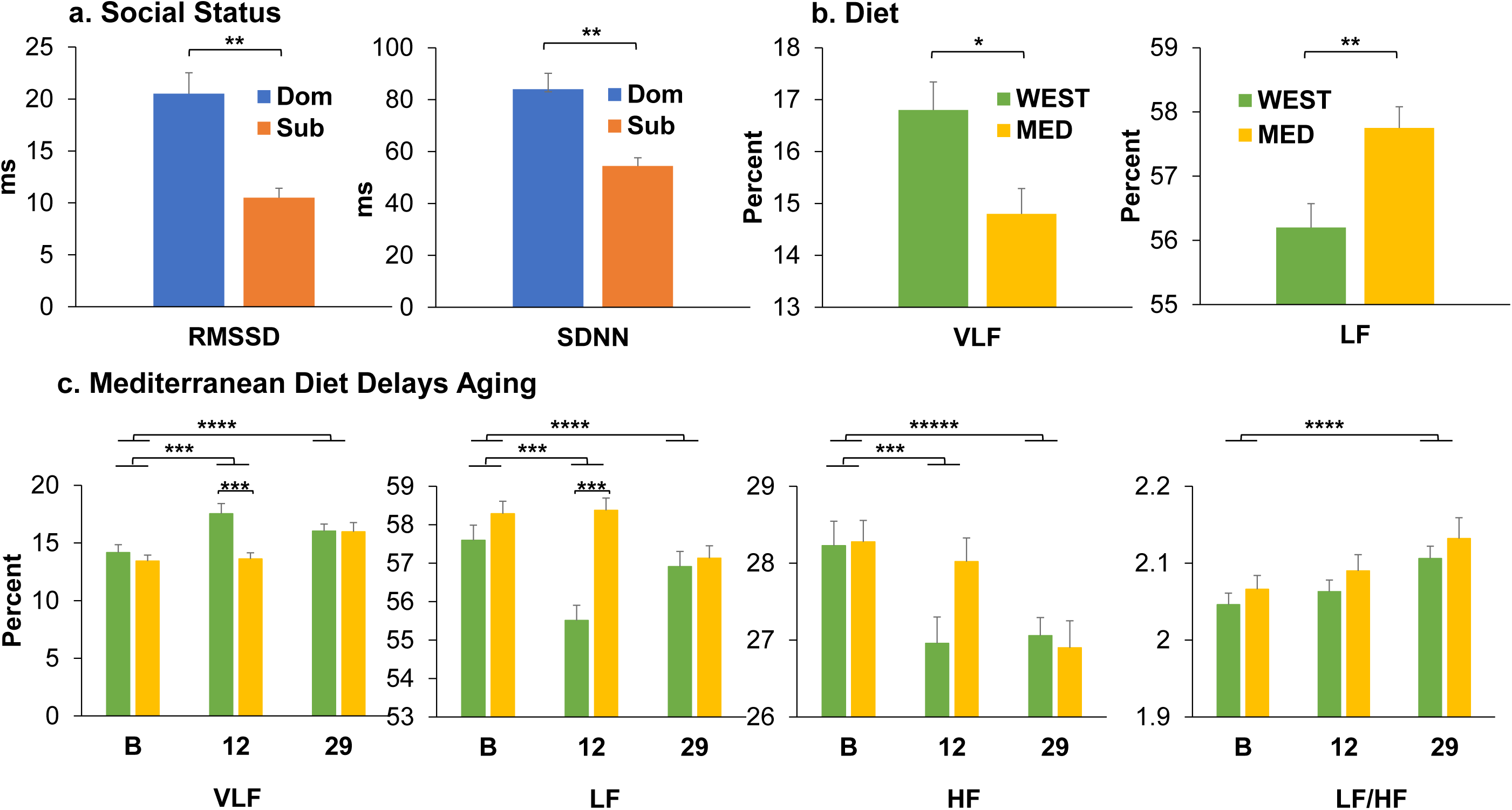
Heart Rate Variability (HRV). HRV was measured at Baseline, 12 months, and 29 months of Western (WEST) or Mediterranean (MED) diet in subordinate (Sub) and dominant (Dom) NHPs. HRV was measured as the standard deviation of successive interbeat intervals from which artifacts have been removed (SDNN), and the root mean square of successive interbeat intervals measured between R-to-R peaks (RMSSD). Fast Fourier transform was used to determine very low frequency (%VLF; range: <0.01Hz), low frequency (%LF; range: 0.01-0.2Hz), high frequency (%HF; range: 0.2-0.8Hz), components, and the ratio of LF/HF. **a. Social Status Differences**. During the Experimental phase dominants had higher RMSSD (F[1, 34]=6.526, p=0.015) and SDNN (F[1, 34]=4.685, p=0.038) than did subordinates. **b. Diet Effects**. The MED group had higher %LF (F[1, 34]=4.322, p=0.045) than the WEST group. %VLF tended to be lower for MED animals, however, the difference was not statistically significant (F[1, 34]=3.453, p=0.072). **c. Changes with Aging and Modification by Diet**. Between Baseline and 12 months, %VLF (F[1, 108]=7.492, p=0.007) increased, whereas %LF (F[1, 108]=5.608, p=0.020) and %HF (F[1, 108]=7.442, p=0.007) decreased. Between Baseline and 29 months, %VLF (F[1, 108]=11.417, p=0.001) and LF/HF (F[1, 108]=12.003, p=0.001) increased, and %LF (F[1, 108]=4.125, p=0.042) and %HF (F[1, 108]=19.413, p<0.0001) decreased. **Age-related changes in %VLF and %LF were delayed by the MED diet** (diet X time interactions: %VLF F[2,179]=4.876; p=0.009; %LF F[2, 179]=5.249, p=0.006). The WEST diet group had significantly higher VLF and lower LF at 12 months (Tukey’s p’s≤0.004), but there was no difference between diet groups at 29 months (Tukey’s p’s>0.10). The same trend was observed in %HF but did not reach significance (diet X time interaction F=2.466, p=0.088). *≤0.10; **≤0.05; ***≤0.01; ****≤0.001; *****≤0.0001.

#### 4.b. Mediterranean Diet Delays Aging Effects on Autonomic Nervous System (ANS) Function (Figure 4c)

Over the course of the experiment, sympathetic activity increased and parasympathetic activity decreased as reflected in power spectrum analyses. Between Baseline and 12 months, and Baseline and 29 months %VLF increased (Baseline versus 12 months: F[1, 108]=7.492, p=0.007; Baseline versus 29 months: F[1, 108]=11.417, p=0.001), whereas decreases were observed in %LF (Baseline versus 12 months: (F[1, 108]=5.608, p=0.020; Baseline vs 29 months: F[1, 108]=4.125, p=0.042) and %HF (Baseline vs 12 months: F[1, 108]=7.442, p=0.007; Baseline versus 29 months: F[1, 108]=19.413, p<0.0001) (**Figure 4c**). In addition, LF/HF increased between Baseline and 29 months (F[1, 108]=12.003, p=0.001) (**Figure 4c**).

The changes over time in %VLF and %LF were delayed by the MED diet (diet X time interactions: %VLF F[2,179]=4.876; p=0.009; %LF F[2, 179]F=5.249, p=0.006). Compared to the MED group, the WEST group had 28.8% higher %VLF (mean difference=3.92) and 4.9% lower %LF (mean difference=2.86) at 12 months (Tukey’s p’s≤0.004), but there was no difference between diet groups at 29 months (Tukey’s p’s>0.10). The same trend was observed in %HF but did not reach significance (diet X time interaction F=2.466, p=0.088).

### 5. Heart Rate Responses to the Acute Stress Test

#### 5.a. Diet and Social Status Effects (Figure 5)

HRs in response to acute stress were examined at 12 and 29 months using ANCOVA controlling for Baseline. At 12 months there were no significant main or interaction effects of diet or social status (all p’s >0.2). At 29 months, there was a significant diet X test time point interaction (**Figure 5a**: F[24, 739]=1.87, p=0.007) suggesting that the two diet groups had different stress response patterns over the course of the test. Specifically, the MED diet group had a greater increase in HR from baseline (MED increase from −30 min to +45 min: MED=60 bpm; WEST= 38 bpm), followed by a decline during recovery not observed in the WEST group (change from +45 min to +240 min: MED= −23 bpm; WEST= + 1.4 bpm). There was also a significant social status X test time point interaction (**Figure 5b**: F[24, 739]=1.58, p=0.038) suggesting that dominants and subordinates had different stress response patterns over the course of the test. Subordinates had a greater and more sustained increase in HR from time 0 than dominants.

**Figure 5.**
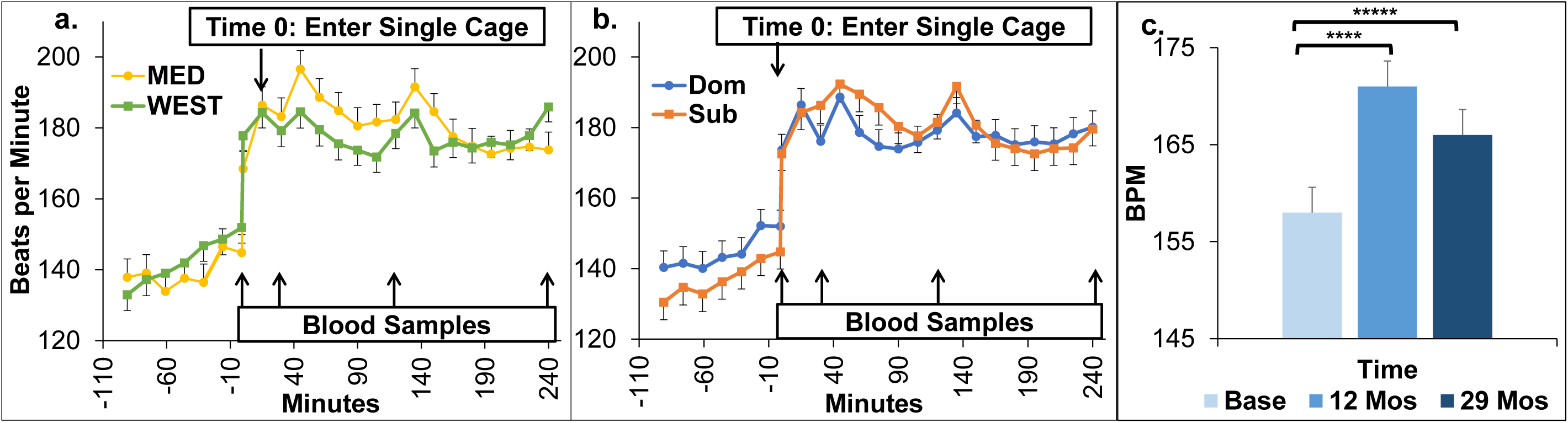
Acute Stress Test Heart Rate. On test day the monkey entered a single cage and a baseline blood sample was taken (Time 0). The caged monkey was moved to an empty room for 30 min. Blood sampling was repeated at 30 min (stress sample) and the monkey was returned to her home pen. Blood samples were taken again at 120 and 240 min (recovery samples) after moving the monkey to a single cage. **Experimental Phase Month 29 Adjusted for Baseline: a. Diet X time interaction** (F[24,739]=1.87, p=0.007): Mediterranean (MED) group had a greater increase in HR from Baseline than the Western (WEST) group; **b. Social status X time interaction** F[24,739]=1.58, p=0.038: Subordinates had a greater and more sustained increase in HR from Baseline than dominants. Adjusted means and SEMs are shown; **c. Heart Rate Increase Over the Course of the Study** F[2, 2411]=120.83, p<0.0001; Baseline vs 12 months Tukey p <0.0001*****; Baseline vs 29 months Tukey p<0.001****.

#### 5.b. Aging Effects (Figure 5c)

HR responses to the acute stress test differed significantly across the three experimental time points (F[2, 2411]=120.83, p<0.0001). Average HRs increased from approximately 158 to 171 beats per minute between the Baseline and 12-month time point (∼8%; Tukey p<0.0001). HR responses then decreased to an average of 166 beats per minute at 29 months (12 vs. 29 months: Tukey p<0.0001), a measure which was still significantly higher than HR responses at Baseline (∼5%; Tukey p<0.0001).

### 6. Cortisol Responses to the Acute Stress Test

#### 6.a. Diet and Social Status Effects

Each experimental time point was examined individually. At 12 months, there was a diet X status interaction (F[1, 34]=4.89, p=0.03). Post hoc comparisons of area under the curves (AUCs) among the 4 groups revealed that dominants in the MED group had lower cortisol responses than dominants and subordinates in the WEST group (Tukey p’s ≤0.01); Supplementary **Figure 2**). This effect was not apparent at 29 months (diet by status interaction: F[1, 33]=0.01, p=0.91; **Supplementary Figure 2**). No other main or interaction effects of social status or diet at any of the three time points reached significance (all p’s>0.11).

#### 6.b. Aging Effects on Cortisol Response to Acute Stress Delayed by MED diet

Visual inspection suggested that the cortisol response changed over the course of the experiment (**Supplementary Figure 2**). To test this, first we compared the time zero (basal) cortisol concentrations between the Baseline Phase, and after 12, and 29 months on diet. There were no significant main or interaction effects of diet or social status (all p’s>0.08), but basal cortisol increased over time (F[2, 68]=37.42, p<0.0001). The 29-month basal cortisol concentrations were significantly higher than those during the Baseline Phase, and after 12 months on diet (Tukey p’s ≤ 0.01; **Supplementary Figure 2**). Next, cortisol AUC was calculated for each of the three experimental time points (Baseline, 12 months, 29 months, **Figure 6a**). ANOVA revealed that the cortisol response to acute stress increased over time (main effect of time: F[2, 68]=23.8, p<0.0001) but that the increase in cortisol was delayed in the MED group (diet X time interaction: F[2, 68]=4.84, p=0.01). To calculate the best estimate of the size of the diet effect, Experimental phase AUCs were evaluated controlling for Baseline AUC with ANCOVA. The cortisol response over the 29-month experimental phase was lower in the MED group than the WEST group (main effect of diet: F[1, 34]=7.33, p=0.011; difference between the adjusted means=2678.2 **Figure 6a**). Compared to the WEST diet, The MED diet significantly reduced the adrenal cortisol response to acute stress by about 15.6%. There were no significant main or interaction effects of social status in any of these analyses (all p’s>0.20).

**Figure 6.**
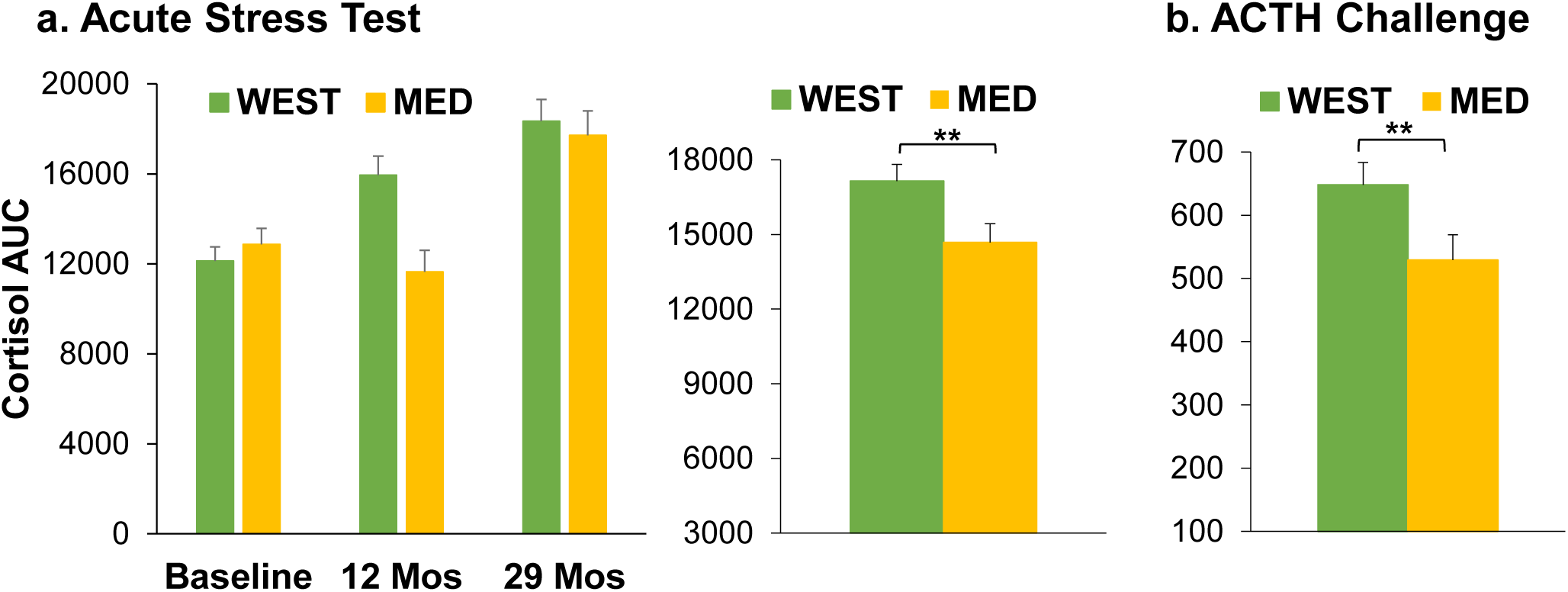
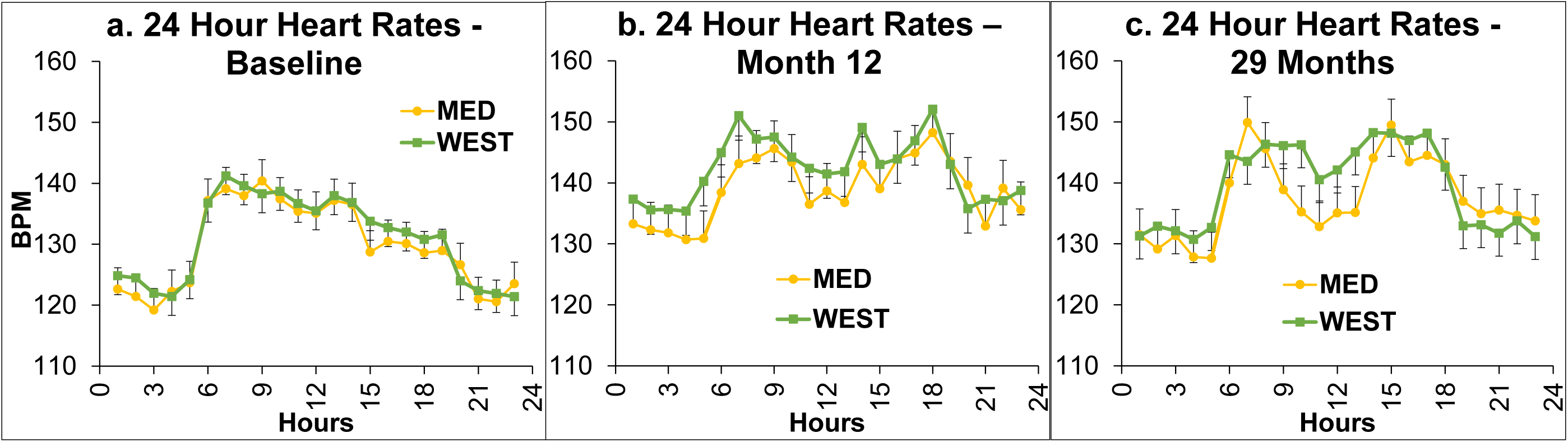
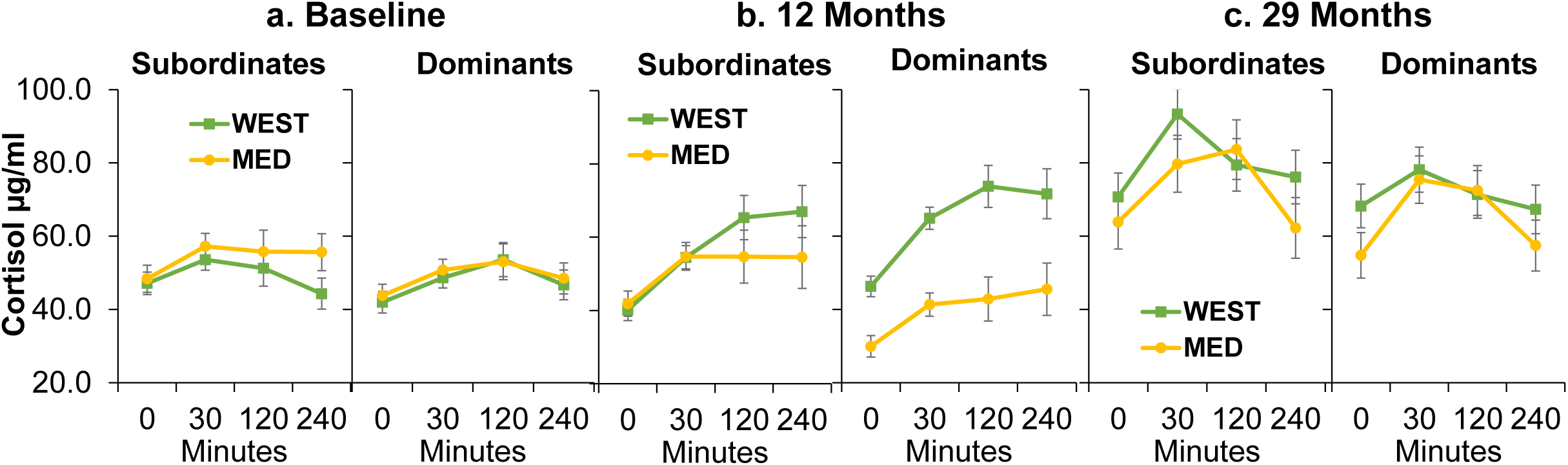
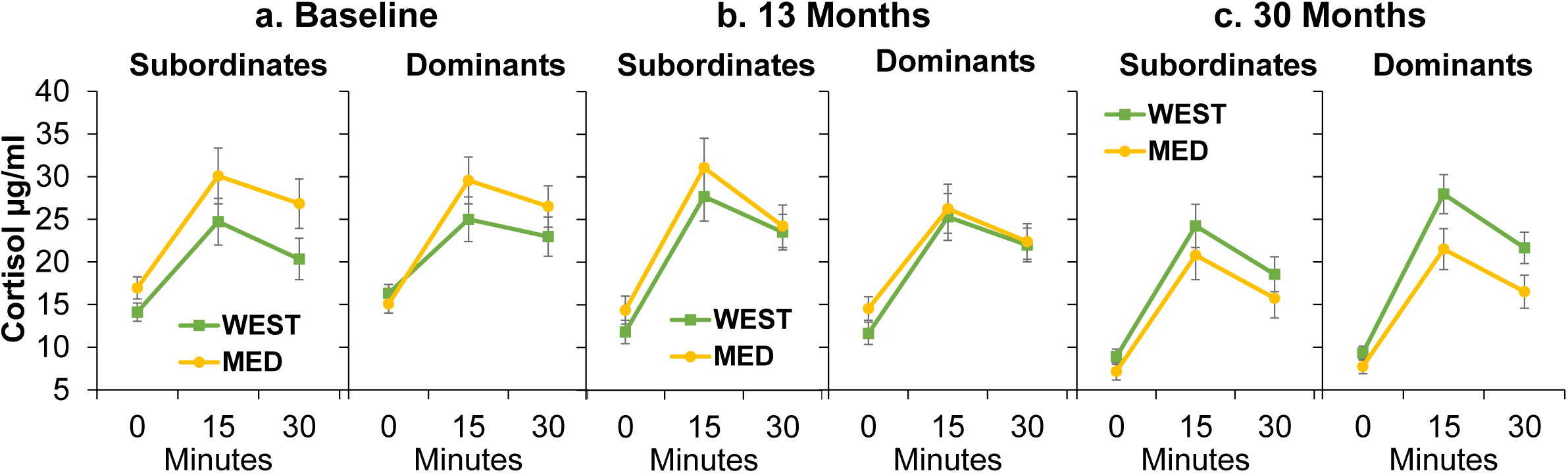
Hypothalamic-Pituitary-Adrenal Function. **a. Cortisol Response to Acute Stress Test**. Cortisol area under the curve were calculated for each of the three experimental time points (Baseline, 12 months, 29 months). **Left:** The cortisol response to acute stress increased over time (main effect of time: F[2, 68]=23.8, p<0.0001) but that the increase in cortisol was delayed by the MED diet (diet X time interaction: F[2, 68]=4.84, p=0.01). **Right:** Experimental phase AUCs were evaluated controlling for Baseline AUC with ANCOVA. The cortisol response over the 29-month experimental phase was lower in the MED group than the WEST group (main effect of diet: F[1, 34]=7.33, p=0.011** (means of the 12 month and 29-month time points adjusted for Baseline are shown). **b. ACTH Challenge Test**. The area under the cortisol curve was calculated for the Baseline and 30-month time points, and effects of status and diet during the Experimental phase were evaluated controlling for Baseline using ANCOVA. The MED diet significantly reduced the adrenal cortisol response to ACTH by about 18% (main effect of Diet: F[1, 33]=9.0, p=0.005**; **Figure 6b**) (means at the 30-month time point adjusted for Baseline are shown).

### 7. ACTH Challenge Test

#### 7.a. Diet and Social Status Effects

Initially the cortisol response to ACTH challenge at the three time points was analyzed separately. There were significant cortisol increases in response to ACTH at all three time points: (Baseline: F[2, 68]=107.6, p<0.0001; 13 months: F[2, 68]=137.9, p<0.0001; 30 months F[2, 66]=25.9, p<0.0001). The cortisol response was higher on average in the MED group at Baseline although this difference did not reach significance (Diet: F[1, 34]=3.4, p=0.07; **Supplementary Figure 3a**). There were no significant effects of social status or diet at 13 months (all p’s>0.10, **Supplementary Figure 3b**). At 30 months those consuming a WEST diet had a greater cortisol response to ACTH than those consuming the MED diet (Diet: F[1, 33]=5.8, p=0.02; **Supplementary Figure 3c**).

The area under the cortisol curve was calculated for the Baseline and 30-month time points, and effects of status and diet during the Experimental phase were evaluated controlling for Baseline using ANCOVA. The MED diet significantly reduced the adrenal cortisol response to ACTH by about 18% (main effect of Diet: F[1, 33]=9.0, p=0.005; difference between the adjusted means=119.3, **Figure 6b**).

## Discussion

This study represents an in-depth interrogation of two main systems, the HPA and the ANS, that are critical for organismal responses and adaptions to changes in the environment. These systems differ in how rapidly they respond to the environment: the ANS has the capacity for a much quicker response and then return to baseline compared to the HPA axis. Consequently, the two systems respond differently to acute and chronic stressors, and each can exert long term effects on health.

This study demonstrates that animals fed a Mediterranean diet have enhanced stress resilience as indicated by lower sympathetic activity, brisker and more overt heart rate responses to acute stress, with more rapid recovery, and lower cortisol responses to acute psychological stress, and to adrenocorticotropin (ACTH) challenge, compared to those consuming a Western diet. Main effects of time also were observed in these middle-aged NHPs. Since the 31-month treatment period is about equivalent to a 9-year follow-up in humans, these time-related changes appear to be due to aging. The Mediterranean diet delayed effects of aging on the ANS and the HPA axis.

### Diet Effects

At mealtime, NHPs moved into single cages on voice command which is accompanied by a cardiovascular response. By study end, the HRs of monkeys consuming the MED diet spiked sharply at mealtimes and recovered significantly in between meals, whereas peaks and recoveries were less pronounced and more sluggish in the WEST group. Likewise, in the acute stress test at study end, the MED diet group had a greater increase in HR from baseline followed by a greater decline during recovery, than the WEST group. Brisk and overt responses to stressors followed by rapid recovery are thought to be adaptive responses that lower disease risk (53, 54). HRV analyses at study end also suggested that the MED group had lower sympathetic activity than the WEST group. Thus, MED diet consumption resulted in stress resilience in the ANS.

Cortisol responses to the acute stress test were higher overall during the Experimental phase in the WEST than the MED group. Likewise, the cortisol response to ACTH challenge was higher in the WEST than the MED diet group, suggesting that the lower cortisol response to acute stress in the MED group may be due in part to lower adrenal responsiveness to ACTH. Thus, MED diet consumption also resulted in stress resilience in the HPA axis. Importantly, compared to the MED group, the WEST diet group ate more, and developed greater adiposity, insulin resistance, hepatosteatosis and different gut microbiome profiles, characteristics which may be related to diet effects on ANS and HPA stress reactivity (38, 39). Future analyses will examine relationships among these dependent variables.

### Social Status Effects

Social status was stable over the 39-month experiment. Subordinates experienced greater social stress than dominants as evidenced by more aggression received, less grooming received, and more time spent fearfully scanning their social group. Subordinates had higher SBP than dominants at Baseline. BP is inversely related to SES and other indices of social disadvantage in humans (55-59), however this is the first report, of which we are aware, of social status differences in BP in NHPs. Subordinates also had lower HRV which also has not been previously reported to our knowledge, and a greater HR rise in response to acute stress. Both of these are congruent with higher SNS:PNS activity, although power spectral analysis revealed no significant differences in the power spectra. It may be that both PNS withdrawal and SNS activation contributed to the HRV differences, with neither component alone reaching significance. Nonetheless, lower HRV is associated with increased cardiovascular and all-cause mortality in humans (60).

Socially-dependent ANS perturbations are important because atherosclerosis, the major pathology underlying coronary heart disease (CHD), is exacerbated by high sympathetic activity, and is more extensive in subordinate than dominant female cynomolgus macaques (61). Likewise, in humans, SES is inversely related to CHD risk (62), and high HR and low HRV also predict CHD (63). However, the relationship between ANS activity and SES is less well understood. Low SES humans have impaired recovery of HRV after acute stress exposure (64). Metabolic syndrome, a powerful risk factor for CHD, is inversely related to SES in humans and to social status in female cynomolgus macaques (25, 65). The human social gradient in metabolic syndrome is explained in part by low frequency power; likewise metabolic syndrome in female cynomolgus macaques is accompanied by high 24 hour HRs (57, 66). Thus, ANS perturbations due to the stress of low social status may have direct effects on the cardiovascular system or work through other risk factors such as metabolic syndrome to increase CHD.

### Aging-Related Changes Over the Course of the Study

This 31-month study was equivalent to a follow up period of about 9 human years. Thus, in these middle-aged monkeys, aging may be a significant factor. All four assessments of ANS function showed uniform changes over time that are likely due to aging. DBP increased in all animals over the course of the study. This observation is congruent with those in women in which loss of parasympathetic tone, and increased sympathetic tone with aging contributed to increased blood pressure (67). Consistent with this observation, twenty-four-hour HRs, and HR responses to the acute stress of social separation increased over the study, and HRV analyses consistently showed that sympathetic activity increased, while parasympathetic activity decreased over the course of the experiment.

Aging in human and nonhuman primates is widely thought to be accompanied by increased cortisol exposure, although, most of the available data are from cross-sectional studies. A range of cortisol concentrations are found in the elderly. Among elders, higher cortisol concentrations are associated with poorer health including greater cognitive impairment, depression, anxiety, frailty and Alzheimer’s disease (68, 69). Basal cortisol and the cortisol response to acute stress also increased with age in this study. These observations are both consistent with increased production and increased insensitivity to cortisol negative feedback. Further longitudinal interrogation of the HPA axis to determine specific functional changes is warranted.

### Mediterranean Diet Delayed Aging Related Changes in ANS and HPA Function

While the SNS:PNS balance shifted toward the SNS over the course of the study as the NHPs aged, this effect was delayed by at least 12 months by the Mediterranean diet. Likewise, the increased cortisol response to acute stress with aging was also delayed by at least 12 months by Mediterranean diet consumption.

### Other Changes Over Time

SBP responses changed differently over time in subordinates and dominants: SBP at Baseline was significantly higher in subordinates than dominants and declined, whereas dominant SBP increased; thus SBP in dominants and subordinates was equivalent by study end. The differences at Baseline suggest that, while BPs were measured 6 months after group formation, subordinates were still stressed by new group formation relative to dominants. Decline over the course of the study suggests that subordinate habituation to their social environment continued over a longer time period. The increase in SBP in dominants over the course of the study may be due to aging.

### Strengths and Limitations of the Study

Major strengths of this study which increase rigor and reproducibility include the experimental design of a controlled, randomized, preclinical NHP trial, confirmation that experimental groups were matched on key Baseline characteristics that might influence outcomes, and the use of Baseline covariates to control for individual differences; practices common in clinical but not preclinical studies. In contrast to human studies, type and quantity of food intake over a long time period was known, and many environmental variables that may affect physiology were controlled including housing, light/dark cycles, and social living characteristics. Three limitations warrant consideration: 1) While Western-like diets have been fed to macaques for several decades, the MED diet formulation was novel and had not been previously fed to NHPs; 2) future investigations are needed to determine whether diet composition affects stress responses in males; and 3) the NHPs studied here have menstrual cycles year round, however tests reported here could not be done with respect to menstrual cycle phase which could influence outcomes.

### Conclusions

These data provide compelling evidence that diet modifies physiological stress responses which has significant implications for human health. Furthermore, while aging shifted ANS function toward increased sympathetic tone, the MED diet resulted in lower sympathetic tone relative to the WEST diet, delaying the effects of aging. Likewise, the MED diet delayed the aging-related increases in cortisol responses to acute stress. Most animal studies are conducted in subjects consuming commercially available chows, designed for species-specific optimal health, that have little resemblance to human diets. This may be an important contributing factor in the frequent failure of preclinical studies to result in interventions that are effective in improving human health. Future preclinical science should consider adopting a dietary background appropriate to the target population.

The observation that diet modifies physiological stress responses is critically important to the public health. Although deleterious effects of psychosocial stress on disease risk are well recognized, efficacious population level interventions to reduce stress responsivity and associated disease have not been identified. However, efficient, population-wide dietary intervention programs are not without precedent. The National Cholesterol Education Program resulted in significant decreases in dietary fat and cholesterol intake, and circulating cholesterol concentrations (70). Likewise, the FDA requirement to list trans fats on food labels and to reduce trans fats in foods resulted in a greater than 50% decrease in circulating levels of these health damaging fats (71) demonstrating the feasibility of population-level diet modification. Based on the findings reported here, the Mediterranean diet pattern may serve as a dietary strategy to reduce the deleterious effects of stress on health without the side effects of medications typically prescribed to manage stress responsivity, and may have a significant public health impact. The Mediterranean diet represents a cost-effective intervention to increase resilience to psychological stress with the potential for widespread efficacy.

## Acknowledgements

We thank the Wake Forest Primate Diet Laboratory for advice on diet formulation and diet preparation, and the Data Management Unit for computer support. We also thank James D. Bottoms, Maryanne Post, Edison Floyd, Nigel Bethel, Terrell Jones, Jenny Estes, Ryan Debo, Kris Michelson, Dewayne Cairnes, Stacey Combs, Naomi Bean, Joshua Long, Jason Lucas, Anna Fimmel, Abigail Beitel, Hannah Register, Jamie Justice, David Neely, Rebecca Marcus, Dorothy Dobbins, and Chrissy May Long for technical assistance.

## Sources of Financial Support

This work was supported by NIH R01HL087103 (CAS), NIH RF1AG058829 (CAS), NIH R01 HL122393 (TCR), U24 DK097748 (TCR), WF ADRC P30AG049638, WF Claude D. Pepper OAIC P30AG012332, and an Intramural Grant from the Department of Pathology, Wake Forest School of Medicine (CAS).

## CRediT Author Statement

**Carol Shively**: Conceptualization, Methodology, Validation, Formal analysis, Investigation, Resources, Data Curation, Writing-Original draft, Visualization, Supervision, Project administration, Funding acquisition. **Susan Appt:** Conceptualization, Methodology, Resources, Writing – Review & Editing. **Haiying Chen:** Formal Analysis, Writing – Review & Editing. **Stephen Day:** Investigation, Writing – Review & Editing. **Brett Frye:** Formal Analysis, Writing – Review & Editing, Visualization. **Hossam Shaltout**: Investigation, Formal Analysis, Writing – Review & Editing. **Marnie Silverstein-Metzler:** Investigation, Writing – Review & Editing. **Noah Snyder-Mackler:** Writing – Review & Editing. **Beth Uberseder:** Methodology, Validation, Data Curation, Investigation, Writing – Review & Editing. **Mara Vitolins:** Conceptualization, Methodology, Writing – Review & Editing. **Thomas Register:** Conceptualization, Methodology, Validation, Investigation, Resources, Writing – Review & Editing.

## Notes

### Competing Interest Statement

The authors have declared no competing interest.

